# Recurrence quantification analysis of heart rate variability is a COVID-safe alternative to gas analysis in the detection of metabolic thresholds

**DOI:** 10.1101/2021.03.22.436405

**Authors:** G. Zimatore, L. Falcioni, M.C. Gallotta, V. Bonavolontà, M. Campanella, M. De Spirito, L. Guidetti, C. Baldari

## Abstract

The first aim of the study was to verify if in individuals with different physical fitness levels the Recurrence Quantification Analysis (RQA) of Heart Rate Variability (HRV) time series could be an alternative to Gas Exchange (GE) analysis in the determination of metabolic thresholds. The second aim was to investigate the validity of the RQA method compared to the GE method in thresholds detection. The two metabolic thresholds were estimated in thirty young individuals during Cardiopulmonary Exercise Testing on a cycle-ergometer and HR, VO_2_ and Workload were measured by the two different methods (RQA and GE methods). RM ANOVA was used to assess main effects of methods and methods-by-groups interaction effects for HR, VO_2_ and Workload at the aerobic (AerT) and the anaerobic (AnT) thresholds. Validity of the RQA at both thresholds was assessed for HR, VO_2_ and Workload by Ordinary Least Products (OLP) regression analysis, Typical Percentage Errors (TE), Intraclass Correlation Coefficients (ICC) and the Bland Altman plots. No methods-by-groups interaction effects were detected for HR, VO_2_ and Workload at the AerT and the AnT. The OLP regression analysis showed that at both thresholds RQA and GE methods had very strong correlations (*r* >0.8) in all variables (HR, VO_2_ and Workload). Slope and intercept values always included the 1 and the 0, respectively. At the AerT the TE ranged from 4.02% to 10.47% (HR and Workload, respectively) and in all variables ICC values were excellent (≥0.85). At the AnT the TE ranged from 2.61% to 6.64% (HR and Workload, respectively) and in all variables ICC values were excellent (≥0.89). Our results suggest that the RQA of HRV time series is a COVID-safe approach for the determination of metabolic thresholds in individuals with different physical fitness levels, therefore, it can be used as a valid method for threshold detection alternative to gas analysis.

## Introduction

Due to the present COVID-19 pandemic, it is currently extremely difficult to perform safely the Cardiopulmonary Exercise Testing (CPET). This gold standard method allows the analysis of oxygen uptake (VO_2_), carbon dioxide production (VCO_2_), and ventilatory measures during an exercise test. In particular, in order to move continuously the flux air from the subject to the device, the modality of gas sampling involves the use of a face mask or a mouthpiece [1]. Therefore, given the difficulty of providing a safe and hygienic environment during every test, subjects and operators may not be appropriately protected from the risk of infection. Thus, it is necessary to develop an additional method as an alternative to Gas Exchange (GE) analysis in order to determine metabolic thresholds during an incremental exercise test. In a previous paper we proposed a new non-linear method based on Recurrence Quantification Analysis (RQA) of Heart Rate Variability (HRV) time series to estimate the aerobic threshold (AerT) in obese subjects [2]. In this special population, the AerT represents a useful parameter to identify the most appropriate physical exercise intensity in order to reduce body weight and to improve physical fitness [3]. An additional metabolic parameter frequently used during exercise testing is the anaerobic threshold (AnT). In fact, after the AerT the body leads to a greater production of lactate with the consecutive rise in ventilation in order to compensate metabolic acidosis. This is considered as an aerobic-anaerobic metabolism phase which ends with a “break-away” in ventilation which appears to correspond to the anaerobic threshold. Therefore, this second threshold represents the onset of hyperventilation and of a predominantly anaerobic exercise phase [4–5]. The AerT and AnT are not always clearly distinguishable in subjects with low level of fitness such as obese individuals. This population cannot perform an incremental physical exercise for long enough and consequently usually the AerT is the only metabolic parameter that can be determined [6]. On the contrary, the AerT and AnT can be more easily observed in a healthy population with a higher fitness level as athletes. Therefore, this population should be considered for developing a new method as an alternative to gas analysis to estimate the metabolic thresholds. Previous studies in athletes proposed several methods to provide information about the relationship between heart rate and the AerT and AnT [7–12]. However, a method based on RQA of HRV time series to distinguish the two metabolic thresholds still needs to be established. Nowadays it is extremely easy to monitor the heart rate using low-cost, non-invasive, and mobile systems [13]. The use of these devices does not necessarily require the presence of the subject in the laboratory, thus, from a practical point of view, the RQA of HRV time series could be an extremely useful method for assessing sport performance and for planning training intensity in athletes in situations where gas analysis is not permitted.

The purpose of the study was to verify if the RQA of HRV time series proposed for obese individuals can be applied also to healthy young subjects. In particular, the first aim was to assess if in individuals with different physical fitness levels the new non-linear method based on RQA could be an alternative to gas analysis in the determination of metabolic thresholds. The second aim of the study was to investigate the validity of the RQA method compared to the GE method in thresholds detection.

## Materials and methods

In this paper the two evaluation methods were called Gas Exchange method (GE) and Recurrence Quantification Analysis method (RQA). In addition, the two thresholds, Aerobic (AerT) and Anaerobic (AnT), were called Aerobic Gas Exchange Threshold (AerT_GE_), Anaerobic Gas Exchange Threshold (AnT_GE_), Aerobic Recurrence Quantification Analysis Threshold (AerT_RQA_), and Anaerobic Recurrence Quantification Analysis Threshold (AnT_RQA_).

### Participants

Thirty subjects (2 females; 28 males) (age = 15.7 ± 2.7 years) were recruited for this study. Participants were divided in three groups: competitive rowers (group A, n=8), recreational rowers (group B, n=8) and other recreational sports (group C, n=14). The A group was made up of competitive rowers, i.e. athletes who performed a minimum of 5 workouts per week and have participated in a regional and/or national competitions in the previous year. The subjects of group B attended rowing activities twice a week. Both the A and B groups were recruited from the Circolo Canottieri Tirrenia Todaro of Rome. The C group was made up of subjects who practice recreational activities different to rowing (twice a week).

All participants underwent clinical examinations to exclude any side effects to physical activity. However, a medical certificate for competitive sport activities or noncompetitive sport activities was requested. Moreover, subjects of all groups had the same main anthropometric characteristics (see Table 1). The exclusion criteria were neuropathy, autonomic dysfunction, cardiovascular diseases. All subjects (or their parents if <18 years) provided a written informed consent before the beginning of the study. This study was conducted in accordance with the Declaration of Helsinki and approved by the CAR-IRB - University of Rome “Foro Italico” Committee (Approval N° CAR 37/2020).

**Table 1.** Characteristics of the study sample (Mean value ± SD)

| Parameters | All (n=31) | A (n=8) | B (n=8) | C (n=14) |
| --- | --- | --- | --- | --- |
| Age (years) | 15.7± 2.7 | 17.13± 4.22 | 16.0± 2.0 | 14.3± 1.6 |
| Height (cm) | 176.6± 8.9 | 183.85± 8.05 | 174.6± 6.7 | 174.3± 8.9 |
| Weight (kg) | 70.2± 13.9 | 73.5± 7.7 | 69.9± 11.7 | 68.6± 17.8 |
| BMI(kg/m<sup>2</sup>) | 22.3± 3.4 | 21.7± 1.3 | 22.9± 3.5 | 22.3± 4.2 |
| FM% | 20.9± 5.9 | 17.6± 3.1 | 21.0± 6.5 | 22.8± 6.2 |

### Procedures

After clinical examination and anthropometric measurements, all subjects performed an incremental exercise test on cycle-ergometer. All subjects were tested in the morning (between 9:00 and 12:00 a.m.), and under similar environmental conditions (temperature 21–22° C; humidity 50–60%). Subjects had their usual breakfast at least 90 minutes before the test session. All tests were performed at the Department of Movement, Human and Health Sciences at the University of Rome “Foro Italico”.

### Anthropometric measurements

The following participants’ anthropometric measurements were assessed: body-weight, height, Body Mass Index (BMI) and percent of fat mass (FM%). Weight and height were measured using a scale and a stadiometer to the nearest 0.1 kg and 0.1 cm, respectively. BMI was calculated as the ratio between weight in kg and the square of height in meters (kg/m^2^). FM% was measured by bioelectrical impedance method (BIA AKERN 101 Anniversary, Pontassieve, FI, Italy).

### Incremental exercise test

The incremental exercise test on the cycle-ergometer was performed with the following protocol: participants started with a 1 minute rest period sitting on the bike, followed by 1 minute of unloaded pedaling (0 Watt). The workload was then increased by 20 Watts/minute for the A group (protocol 1) and by 15 Watts/minute for the B and C group (protocol 2), in order to maintain the total exercise time within about 15 minutes. Participants were asked to keep a cadence of 60-70 revolutions per minute (rpm). During the test, perception of physical exertion was assessed using OMNI Scale of Perceived Exertion (0-10 scale, [14]) 15 seconds prior to the end of every stage. The test ended when one of the following conditions was reached: a value of 10 on OMNI Scale of Perceived Exertion, the 90% of the subject’s predicted HRmax (beats/min) or a respiratory exchange ratio equal to 1.1. The HR was recorded by a chest belt (HRM-Dual™, Garmin^®^) contemporarily to Oxygen consumption (VO_2_, mL/min), carbon dioxide production (VCO_2_, mL/min), and pulmonary ventilation (VE, mL/min) that were measured by an automatic gas analyzer (Quark RMR-CPET Cosmed™, Rome, Italy) [15].

### Detection of the AerT and AnT (GE method)

Gas exchange method (GE) is the gold standard to detect both AerT and AnT as reported by Meyer et al. (2005) [5]. The AerT_GE_ (that occurred at time named T_1_), was determined offline for each subject by plotting the ventilatory equivalent of oxygen (VE/VO_2_) as a function of VO_2_ to identify the point where the VE/VO_2_ reached its lowest value during the exercise test [16–18]. This procedure was confirmed by a graph with VCO_2_ on the y axis and VO_2_ on the x axis: two regression lines are fitted for the upper and the lower part of the relation and their intersection represents the AerT_GE_ (V-slope method, [5]).

Similarly, the AnT_GE_ (that occurred at time named T2) was determined by plotting the ventilatory equivalent of carbon dioxide (VE/VCO_2_) as a function of VO_2_ to identify the point where the VE/VCO_2_ reached its lowest value during the incremental test. This procedure was confirmed by a graph with VE on the y axis and VCO_2_ on the x axis: two regression lines are fitted for the upper and the lower part of the relation and their intersection represents the AnT_GE_ [5].

HR e VO_2_ values corresponding at the two thresholds were reckoned as mean values of the last 30 seconds of the workload.

### Recurrence quantification analysis method (RQA)

RQA can be defined as a graphical, statistical and analytical tool for the study of nonlinear dynamical system [19] and it is successfully used in a multitude of different disciplines from physiology [20–21], to earth science [22] and economics [23–24]. This method is widely explained in a previous paper [2]: it was observed that by recurrence analysis, in epoch-by-epoch (with sliding windows) mode (RQE), the rapid shifts from high to low (and vice versa) of percent of determinism (DET, the percentage of recurrence points which form diagonal lines) is usually an indicator of regime changes and phase transitions [25–26]. In fact, DET (High DET means high autocorrelation) is a universal marker of crisis in fields ranging from physiology to finance [27]. Laminarity (LAM, the percentage of recurrence points which form vertical lines), analogous to DET, measures the number of recurrence points which form vertical lines and indicates the amount of laminar phases (intermittency) in the system studied. In the present paper we use both ENT and LAM to study the increase in correlation and to identify the metabolic thresholds; the present study was focused on chaos-order transitions and physical fatigue was considered as an order parameter acting at physiological level. More details can be found at http://www.recurrence-plot.tk/ and in [26, 28–30].

### Data preprocessing

As previously explained by Zimatore et al. (2020) [2], since the heart rate (HR) was continuously recorded and collected breath by breath, the time series of RR interval (temporal variation between the sequences of consecutive heartbeats) was reckoned by the ratio 60000/HR. Moreover, since HR physiologically increases in relation to the workload, this physiological trend was removed in order to analyze the cardiac regime changes (detrended RR interval) for each subject (the theoretical value of best linear fit on original time series were subtracted to every point belong to original time series). The optimization procedure of input parameters was discussed in a previous work [2], where the following same values were used:

- delay (lag) is set to 1,
- embedding dimension is set to 7,
- cut-off distance (radius) to 50% of mean distance between all pairs of points in time,
- line (minimal number of consecutive recurrences to score a determinism line) set to 4.

RQA is computed across consecutive distance matrices corresponding to consecutive and overlapping sliding windows (epochs) along the series, and this mode of analysis is called RQE (Recurrence Quantification by Epochs). The occurrence of an abrupt change of DET and LAM in adjacent windows corresponds to a transition in the dynamical regime of the signal. RQE analysis was carried out by adopting the same parameters setting as for the global mode (in the present study, when an epoch coincides with all points of the time series, the result is called RQA), plus the definition of windows having a length of 100 points (1 point correspond to 1 breath) and shifting of 1 point between consecutive windows, respectively.

RQA software, RQE.exe and RQC.exe version 8.1, are those included in RQA 14.1 [28]. RQE on sliding windows partially overlapped are carried on a detrended RR interval and then RQA measures are obtained. The software used for the RQA analysis, that generates Recurrence plots (RP) and other RQA utilities, is available from http://cwebber.sites.luc.edu.

### Detection of the AerT and AnT (RQA method)

The method of threshold detection based on RQA of RR intervals (HRV time series) can be described as follows: specifically, AerT_RQA_ was determined by the time point when the statistically relevant minimum of percent of determinism and laminarity occurred. AnT_RQA_, instead, occurred when percent of determinism is saturated (DET>90%); both threshold time points correspond to evident changes in RP texture (see Fig 1). HR e VO_2_ values corresponding to the two thresholds were reckoned as mean values on the last 10 values of the original time series before the value corresponding to the two threshold times (T1 and T2).

**Figure 1.**
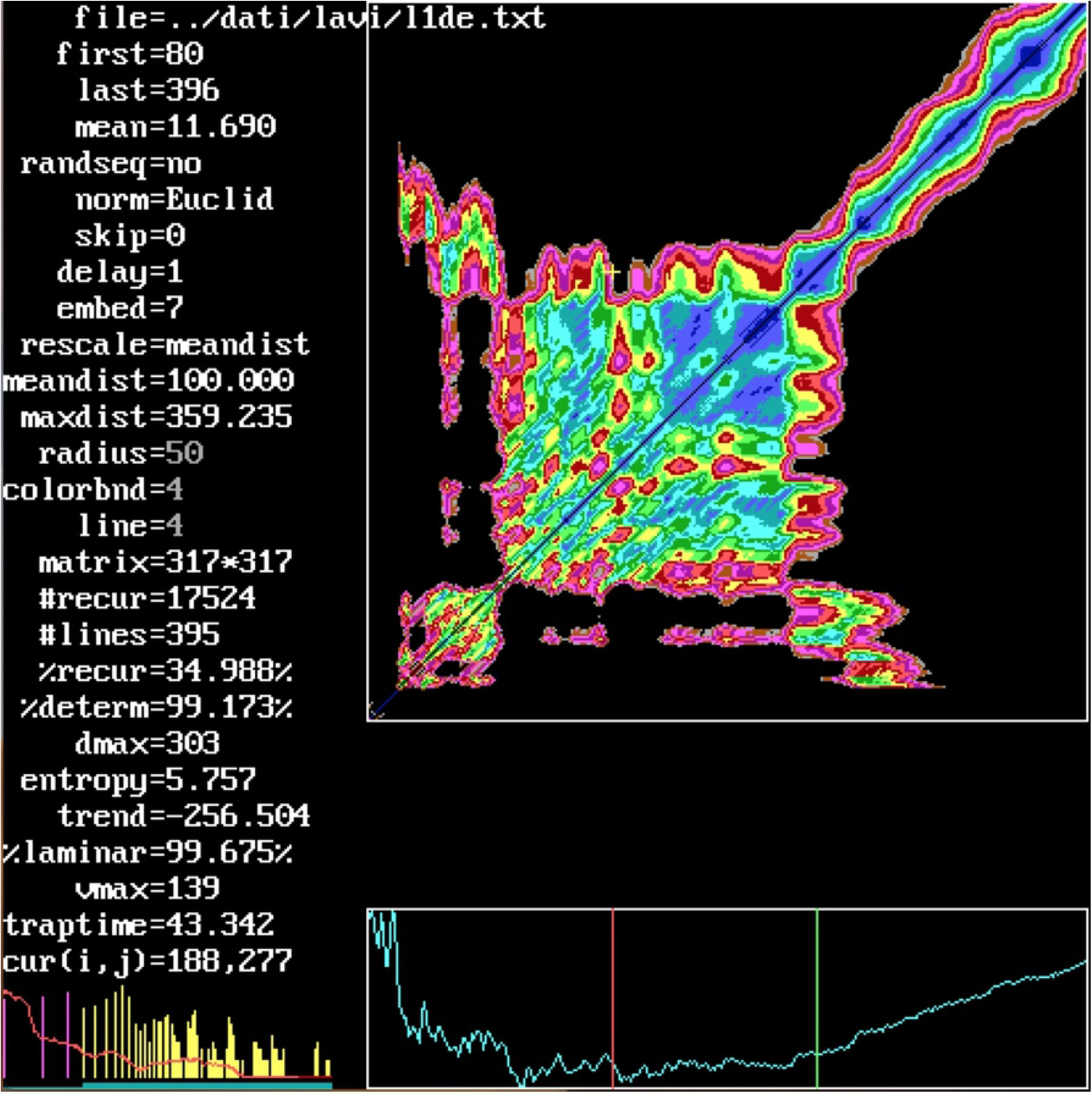
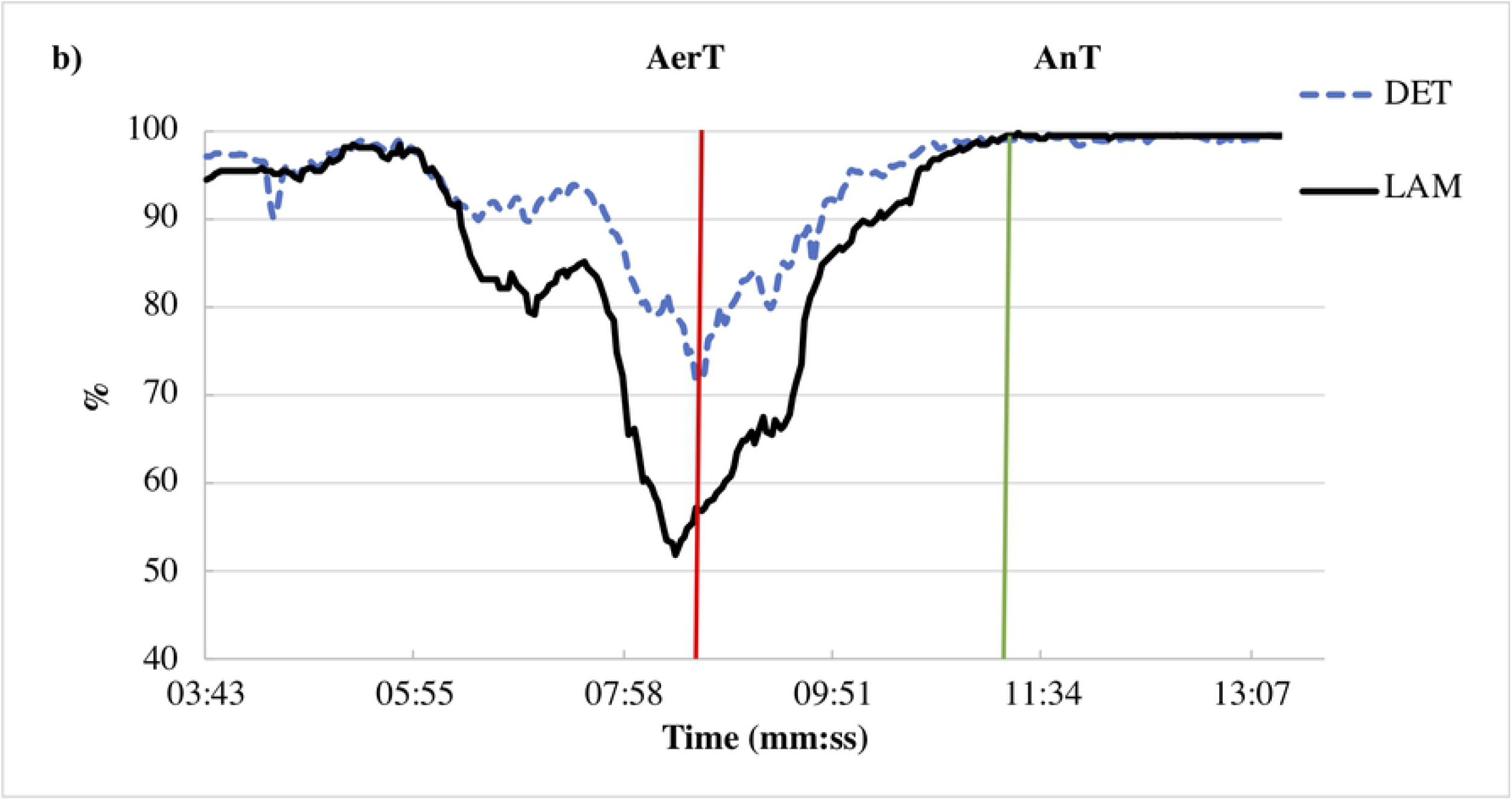
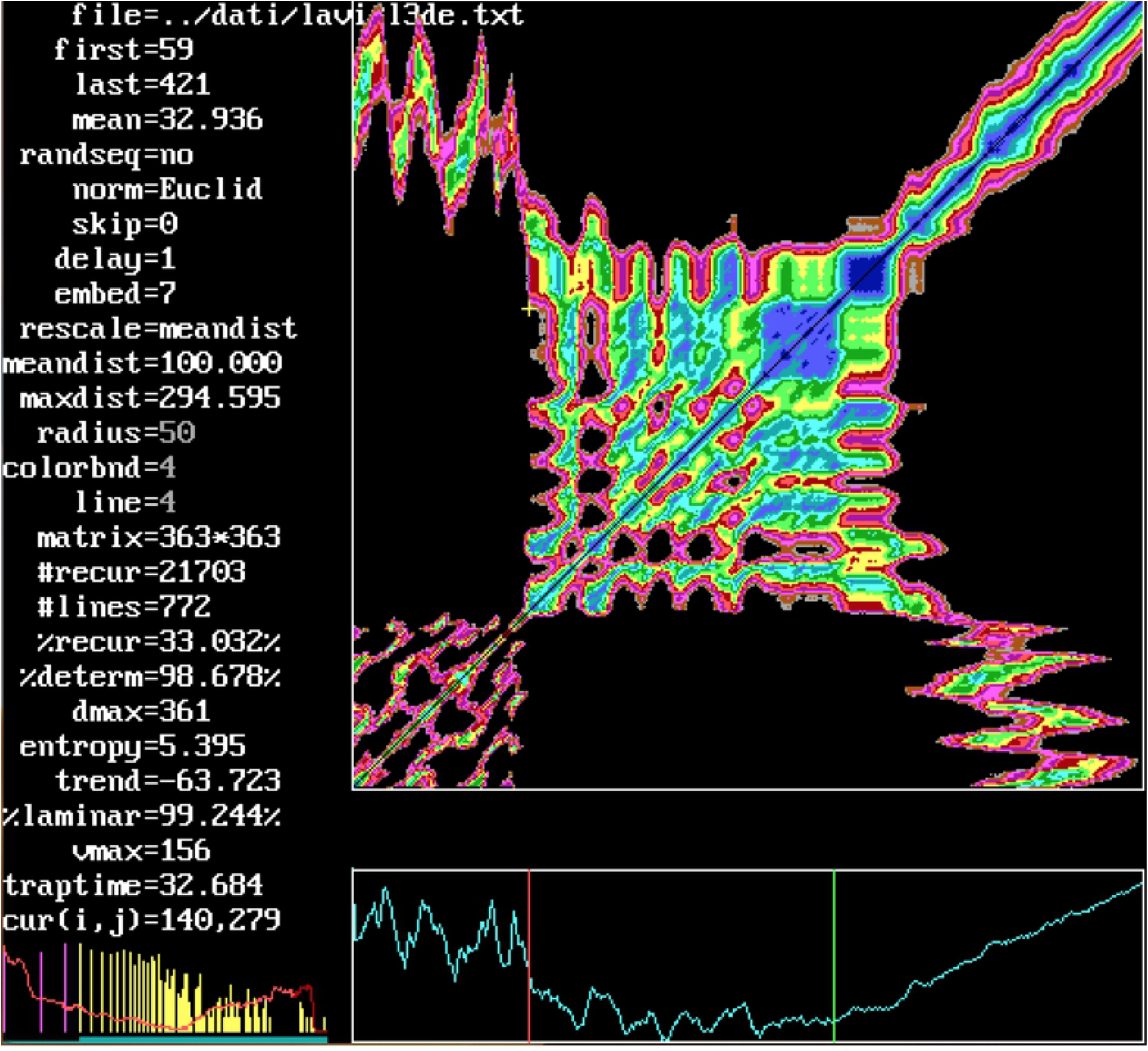
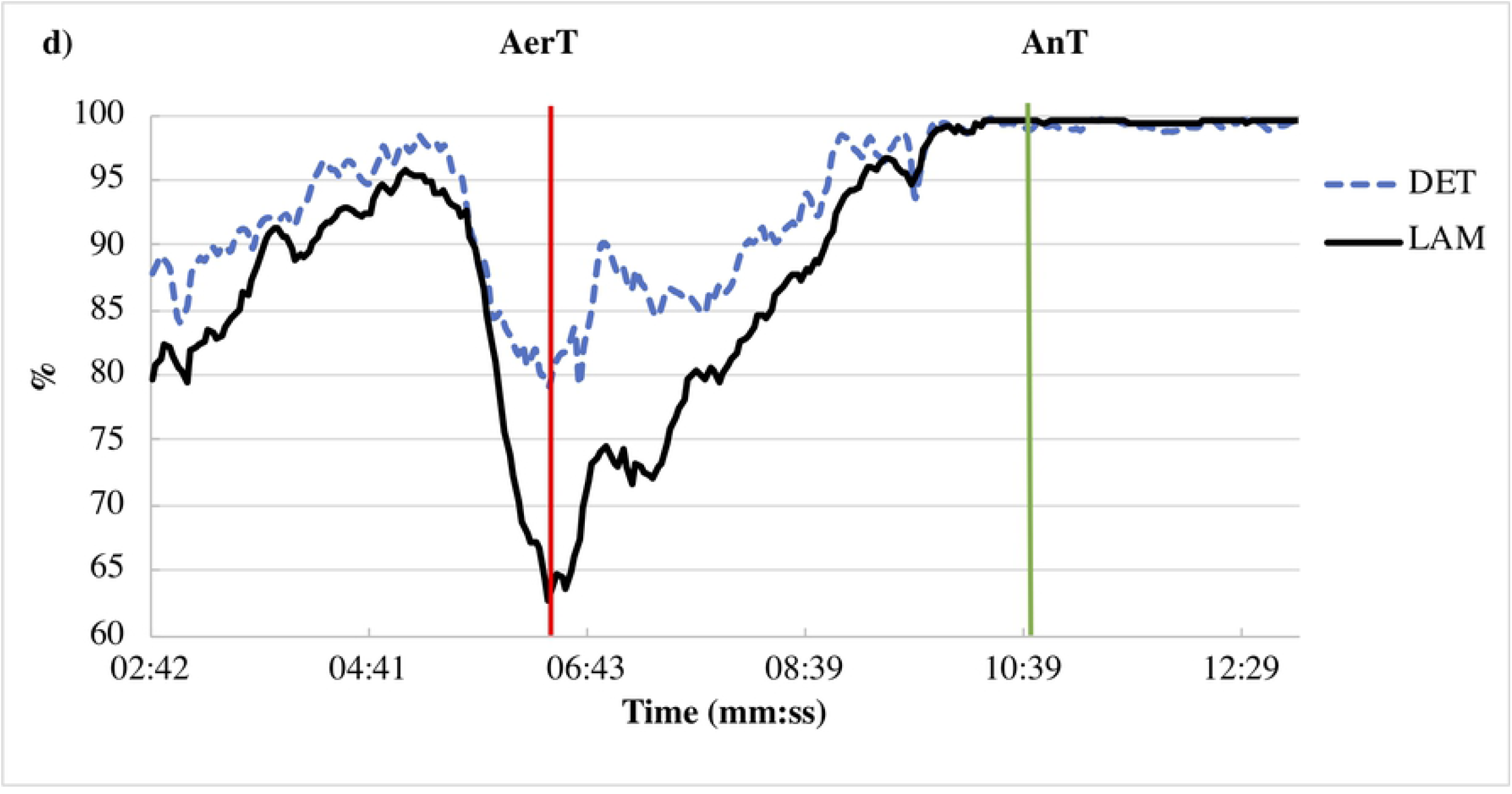
RP and RQE comparison. A comparison among two representative RP is shown in panel a (subject #1) and panel c (subject #3). Panel b (subject #1) and panel d (subject #3) show the percentage of recurrence points which form diagonal lines (DET) and the percentage of recurrence points which form vertical lines (LAM) epoch by epoch vs time (min:sec) of detrended RR interval. The red line enlightens the statistically relevant minimum that corresponds to the AerT and the green line corresponds to the AnT where the percent of determinism reaches saturation. Subject #1 is a competitive rower (group A, protocol 1) and subject #3 is a recreational rower (group B, protocol 2). The test starts at 00:00; AerT occurred at time T_1_, 8:38 and 5:49 for #1 and #3, respectively; AnT occurred at time T_2_, 11:24 and 11:01 for #1 and #3, respectively. A square pattern in the middle is easily distinguishable, the vertical red line is manually placed nearby before the first metabolic transition, the green one nearby before the second one. The colors of RP reflect different Euclidean distances between trajectories and are analogous to geographical relief maps going from blue to red at increasing Euclidean distance. Distances greater than cut-off correspond to black regions.

### Statistical analysis

Repeated measures analysis of variance (RM ANOVA) was used to assess main effects of methods (RQA vs GE) and methods-by-groups (A, B and C) interaction effects for HR, VO_2_ and Workload at the AerT and the AnT. Agreements between RQA and GE methods at the AerT and the AnT were assessed for HR, VO_2_ and Workload by Ordinary Least Products (OLP) regression analysis [31]. Coefficients of determination (R^2^) and regression parameters (slope and intercept) with the 95% of confidence intervals (95% CI) were calculated for the OLP regression equation to determine fixed and proportional biases. The hypothesis of proportional and fixed bias was rejected when the 95% CI contained the value 1 for the slope and the 0 for the intercept, respectively. Percentage differences between the RQA and GE methods at the AerT and the AnT (reported as mean and range values) were calculated for HR, VO_2_ and Workload. RQA method validity was also assessed by comparing HR, VO_2_ and Workload at the AerT and the AnT vs the same variables measured by the GE method with a paired samples t-test. Measurement error was expressed in Typical percentage Error (TE) which was calculated by dividing the standard deviation of the difference percentage by 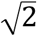. Intraclass correlation coefficients (ICC) was used as a parameter for criterion validity of the RQA method compared to the GE method at the AerT and the AnT for all variables (HR, VO_2_ and Workload). Bland-Altman plots were applied to determine the 95% CI between the RQA and GE methods at the AerT and the AnT for the HR [32].

Statistical significance was defined as p ≤0.05. All statistical analysis was performed by SPSS version 24.0 software (SPSS Inc., Chicago, IL).

## Results

Workload at the AerT and HR, VO_2_, and Workload at the AnT were not statistically different between RQA and GE methods. Significant differences were found between the two methods for HR at the AerT (HR at the AerT_GE_=137.77 ± 15.25 beats/min; HR at the AerT_RQA_=141.44 ± 15.35 beats/min) and for VO_2_ at the AerT (VO_2_ at the AerT_GE_=1849.56 ± 477.68 mL/min; VO_2_ at the AerT_RQA_=1945.37 ± 516.91 mL/min). No methods-by-groups interaction effects were detected for HR, VO_2_ and Workload at the AerT and the AnT (Table 2).

**Table 2.** Main effects of methods (RQA vs GE) and method-by-group (A, B and C) interaction effects in HR, VO2 and Workload at AerT and AnT (ANOVA RM)

| Parameter | Effect | AerT | AnT |
| --- | --- | --- | --- |
| HR (beats(min) | Method | 0.026* | 0.466 |
|  | Method-by-group | 0.766 | 0.789 |
| VO<sub>2</sub> (mL/min) | Method | 0.020* | 0.334 |
|  | Method-by-group | 0.717 | 0.746 |
| Workload (Watt) | Method | 0.115 | 0.288 |
|  | Method -by-group | 0.261 | 0.765 |

\*p<0.05

### HR, VO_2_ and Workload at AerT

The agreements between HR, VO_2_ and Workload at the AerT_GE_ and the AerT_RQA_ are presented in Table 3. In all variables (HR, VO_2_ and Workload) OLP regression analysis showed that the AerT_GE_ and the AerT_RQA_ had very strong correlations (*r* >0.8). Slope and intercept values always included the 1 and the 0, respectively. Mean percentage differences ranged from 2.67% to 5.18% (HR and VO_2_, respectively). HR and VO_2_ at the AerT resulted statistically different between the two evaluation methods (*p* <0.05) while no differences were assessed for Workload at the AerT. The TE for HR and VO_2_ at the AerT was considered acceptable (4.02% and 8.40%, respectively), moreover the TE for the Workload at AerT (10.47%) was slightly higher than the acceptable 10% limit [33]. ICC values were excellent (≥ 0.85) for all variables at the AerT [34–35]. The OLP regression analysis plots of HR, VO_2_ and Workload values at the AerT_GE_ and at the AerT_RQA_ are graphically shown in Figs 2a, 3a and 4a, respectively. In each graph it is described the OLP regression plot with the linear regression (solid line), the identity (dashed line), the equation, the correlation coefficient (*r*) and the absolute mean differences. For the HR the Bland-Altman plot with the mean difference in the solid line (3.7 beats/min) and the 95% CI in the dashed lines (−1.5 to 18.9 beats/min) is shown in the upper-left panel of Fig 2a.

**Table 3.** Agreement between HR, VO_2_ and Workload values at AerT and AnT estimated by GE and RQA methods.

| Parameter (threshold) | R<sup>2</sup> | Slope<br>(95% CI) | Intercept<br>(95% CI) | Mean diff (%)<br>(95% CI) | *p* | TE (%) | ICC<br>(95% CI) |
| --- | --- | --- | --- | --- | --- | --- | --- |
| HR (AerT) | 0.760 | 1.01<br>(0.82 to 1.23) | 2.83<br>(-27.74 to 27.87) | 2.83<br>(0.80 to 4.87) | 0.014* | 4.02 | 0.85<br>(0.68 to 0.93) |
| VO<sub>2</sub> (AerT) | 0.846 | 1.08<br>(0.92 to 1.27) | -56.09<br>(-404.00 to 240.30) | 5.59<br>(1.34 to 9.84) | 0.015* | 8.40 | 0.90<br>(0.78 to 0.96) |
| Workload (AerT) | 0.908 | 1.03<br>(0.91 to 1.17) | 1.30<br>(-11.411 to 12.523) | 5.26<br>(-0.04 to 10.55) | 0.080 | 10.47 | 0.95<br>(0.89 to 0.98) |
| HR (AnT) | 0.792 | 0.96<br>(0.80 to 1.16) | 5.51<br>(-26.62 to 32.19) | -0.33<br>(-1.65 to 0.99) | 0.524 | 2.61 | 0.89<br>(0.78 to 0.95) |
| VO<sub>2</sub> (AnT) | 0.941 | 0.914<br>(0.83 to 1.010) | 183.73<br>(-54.17 to 399.02) | -0.83<br>(-3.24 to 1.59) | 0.319 | 4.78 | 0.97<br>(0.93 to 0.98) |
| Workload (AnT) | 0.947 | 0.94<br>(0.86 to 1.03) | 11.09<br>(-2.42 to 23.40) | 2.82<br>(-0.54 to 6.18) | 0.225 | 6.64 | 0.97<br>(0.94 to 0.99) |

Determinant coefficient ( $R^2$ ), slope and intercept of the regression equations, mean percentage differences (mean diff%), p values, typical percentage error (TE), confidence interval (IC) and intra-class correlation coefficient (ICC). \* $p < 0.05$

**Fig 2.**
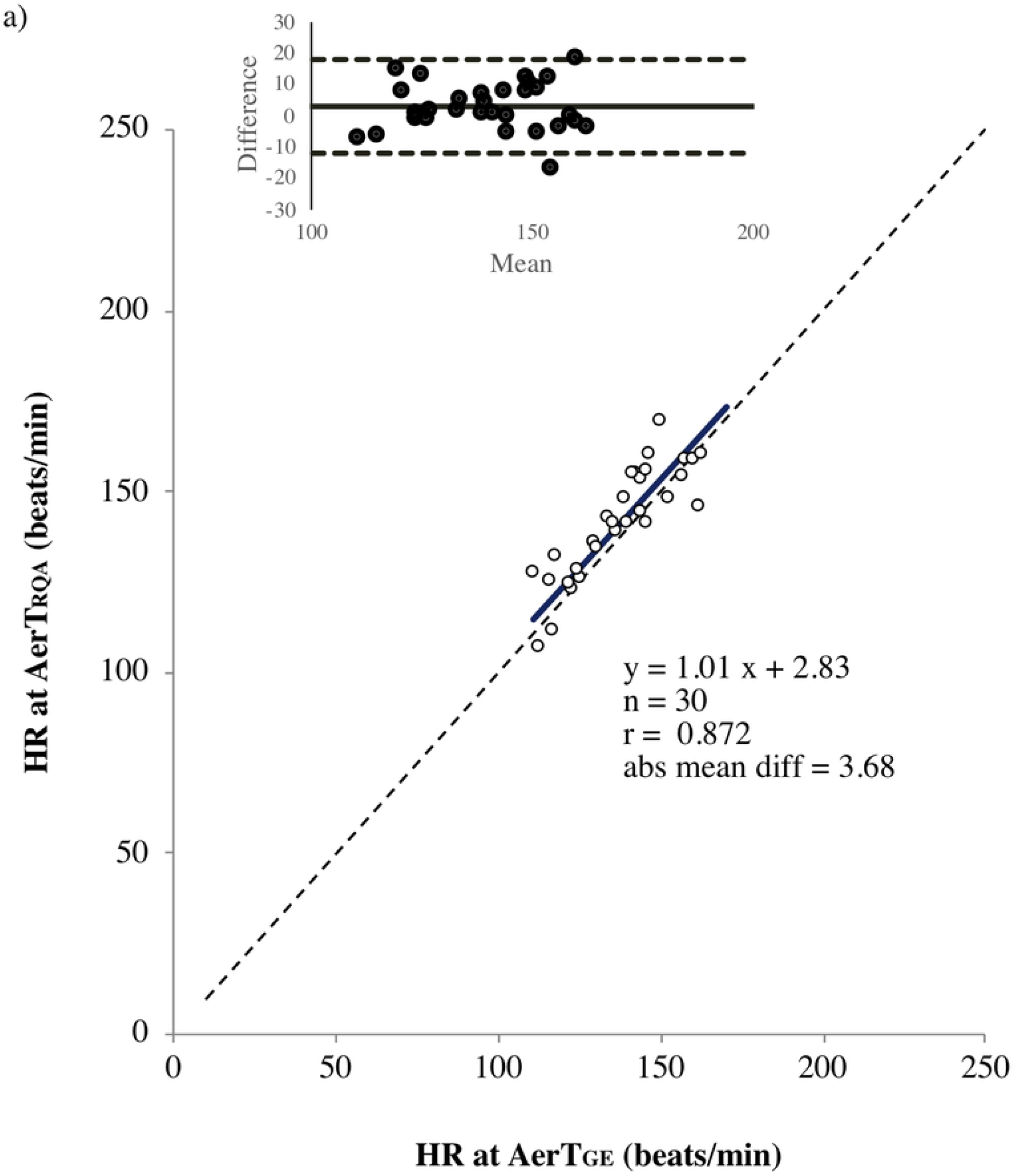
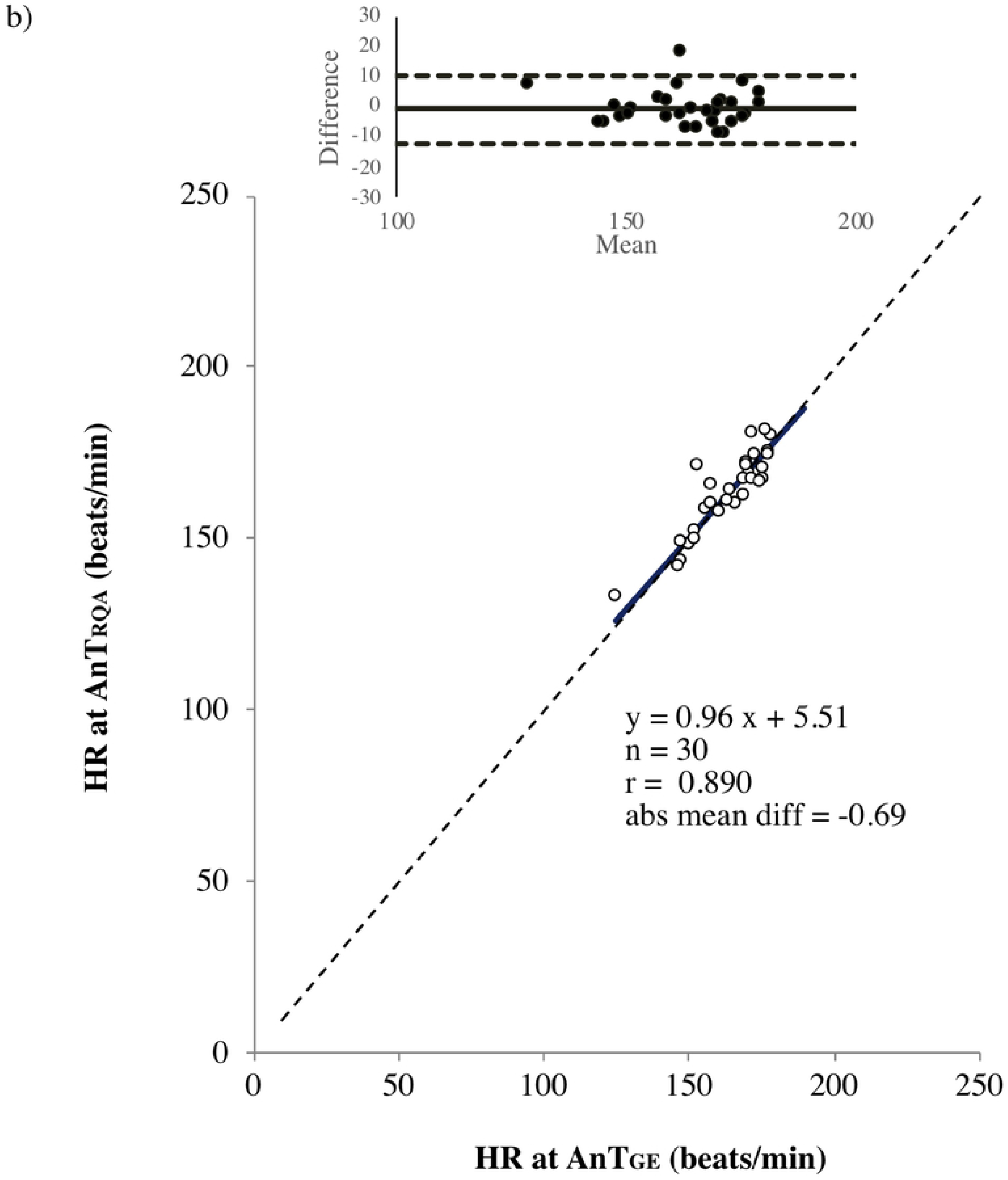
Regression and difference (Bland-Altman) plots of HR estimated by RQA and GE methods at AerT (a) and AnT (b).

**Fig 3.**
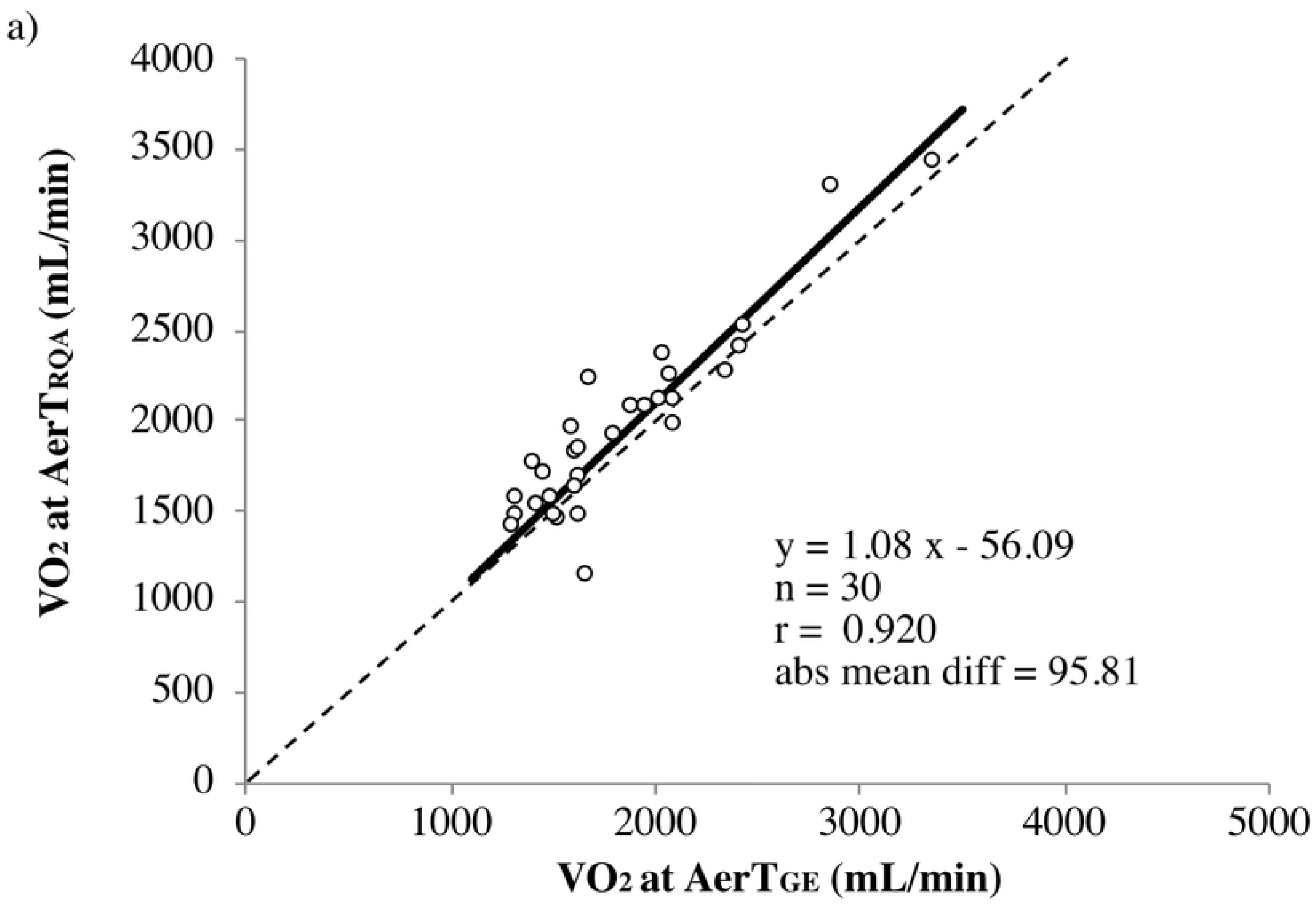
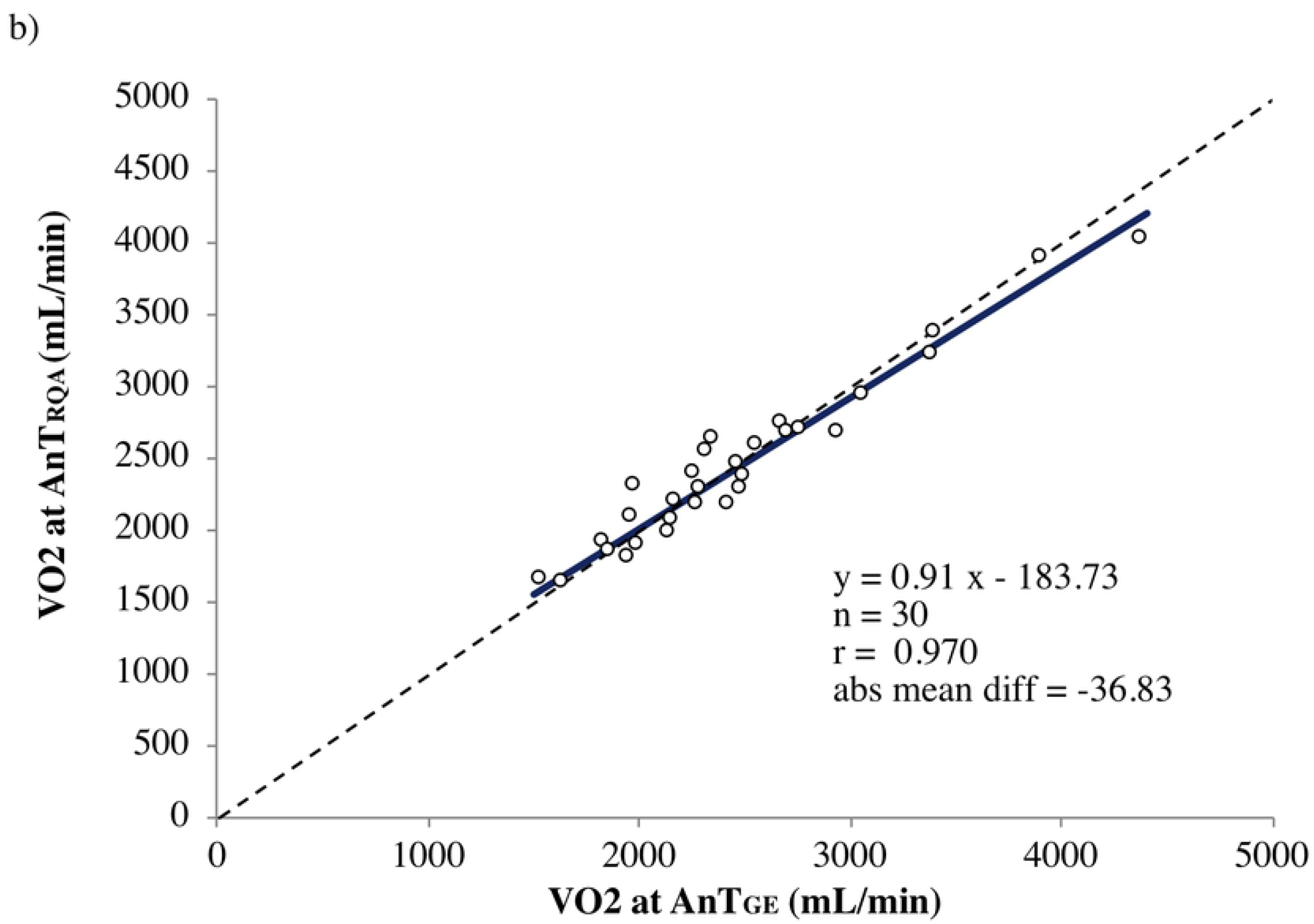
Regression plots of VO_2_ estimated by RQA and GE methods at AerT (a) and AnT (b).

**Fig 4.**
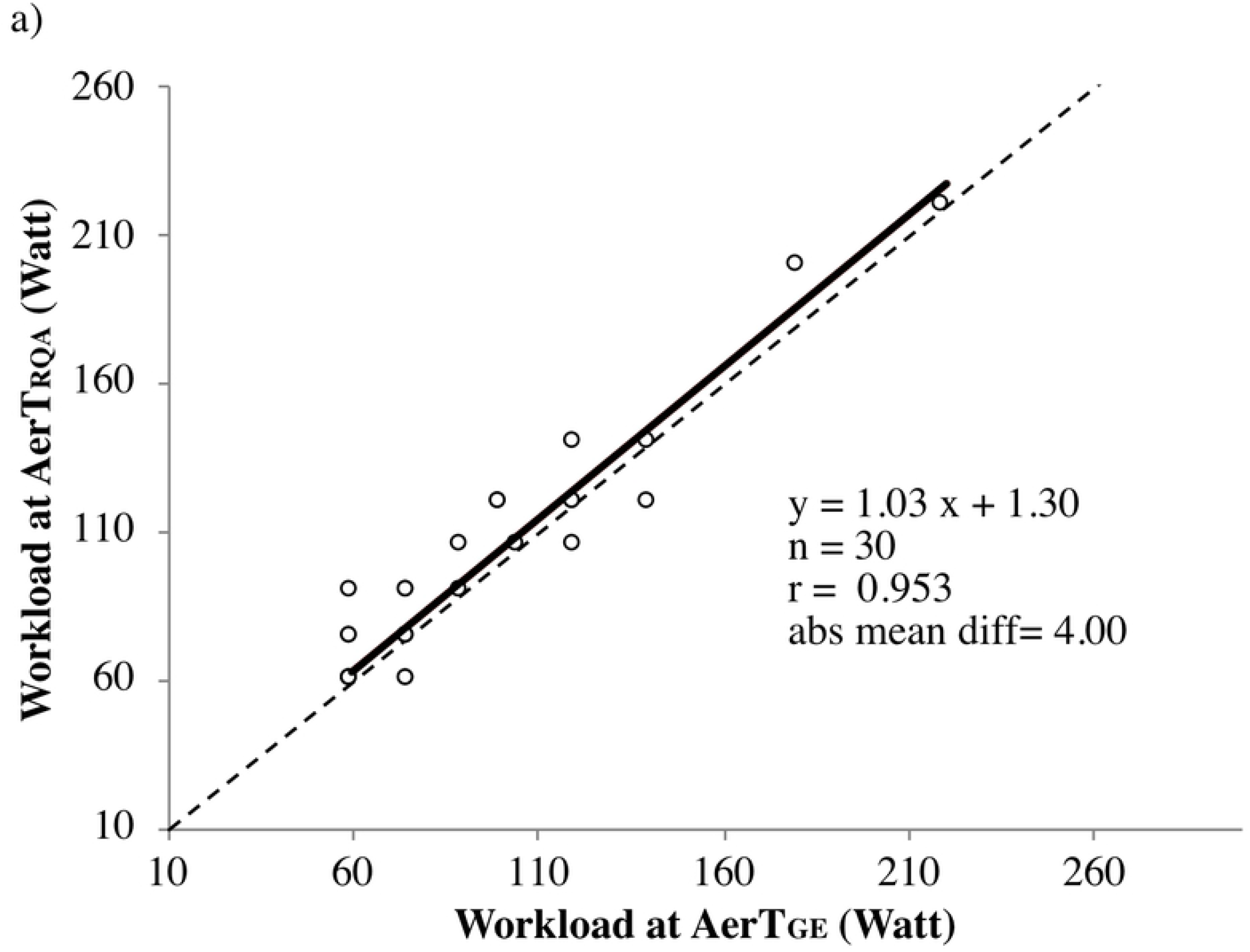
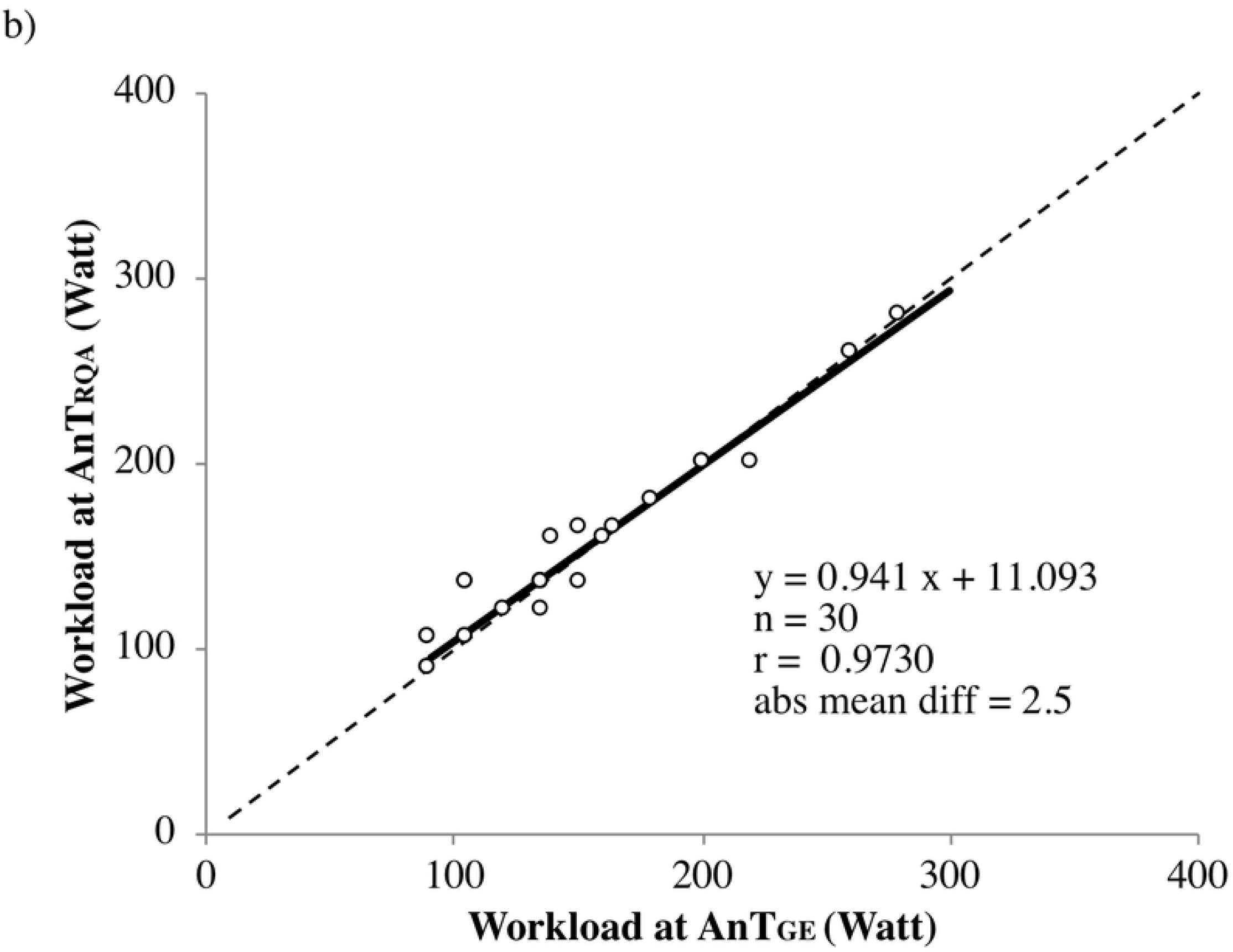
Regression plots of Workload estimated by RQA and GE methods at AerT (a) and AnT (b).

### HR, VO_2_ and Workload at AnT

The agreement between HR, VO_2_ and Workload at the AnT_GE_ and the AnT_RQA_ are presented in Table 3. In all variables (HR, VO_2_ and Workload) OLP regression analysis showed that the AnT_GE_ and the AnT_RQA_ had very strong correlations (*r* >0.8) [36]. Slope and intercept values always included the 1 and the 0, respectively. Mean percentage differences ranged from −0.42% to 1.72% (HR and Workload, respectively) and all variables corresponding to the AnT did not result statistically different between the two evaluation methods. For HR, VO_2_ and Workload at the AnT the TE was considered acceptable (2.61%, 4.78% and 6.64%, respectively) and ICC values were excellent (≥0.89) [33–35]. The OLP regression analysis plots of HR, VO_2_ and Workload at the AnT_GE_ and at the AnT_RQA_ are graphically shown in Figs 2b, 3b and 4b, respectively. For the HR the Bland-Altman plot with the mean difference (−0.7) and the 95% CI (−12.1 to 10.7 beats/min) is shown in the upper-left panel of Fig 2b.

## Discussion

The first aim of the study was to verify if in individuals with different physical fitness levels the new non-linear method based on RQA could be an alternative to gas analysis in the determination of metabolic thresholds. The second aim was to verify the validity of the RQA method compared to the GE method in thresholds detection. To our knowledge, this is the first study assessing the accuracy of the RQA method during the determination of both the AerT and the AnT. In particular, in the present investigation we examined the RQA epoch-by-epoch procedure in order to analyze HRV time series during phase transition. The RQA method is a graphical and analytical approach which has been successfully used in a multitude of different fields as earth science [22], economics [23–24] and physiological signals as otoacoustic emissions [20–21]. However, this methodology still needs to be deeply investigated in the field of exercise testing.

Even though for HR and VO_2_ at the AerT significant differences were found between the two methodologies (RQA and GE methods), no differences were detected for Workload at the AerT and for HR, VO_2_ and Workload at the AnT. Furthermore, the absence of methods-by-group interactions in all variables at both the thresholds indicated that the RQA method is not influenced by the level of physical fitness of the subjects. Therefore, this method can be applied in individuals practicing different sports and with different physical fitness levels in situations where gas analysis is not available. These results permitted us to move to the second aim of the study which consisted in verifying the validity of the RQA method against the GE method during thresholds identification through different validation parameters (OLP regression, TE, ICC, and the Bland Altman plots) individually for HR, VO_2_ and Workload variables corresponding at the AerT and the AnT.

Regarding the degree of agreement between the two methods of AerT detection (AerT_GE_ and AerT_R_QA), all variables (HR, VO_2_ and Workload) showed very strong correlations [36]. Neither fixed or proportional biases were present in all parameters and mean percentage differences were less than 5% for HR and Workload and less than 6% for VO_2_. As observed in the RM ANOVA results, HR and VO_2_ at the AerT showed significant differences between the two methods. However, both these parameters showed an acceptable TE (HR 4.02%; VO_2_ 8.40%) and excellent ICC values (HR 0.85; VO_2_ 0.90) [33–35]. Therefore, for all these reasons the statistical differences detected between the two methods can be considered not relevant for practical applications. In particular, the low percentage of TE in the heart rate indicated that this variable at the AerT_RQA_ represents the most accurate parameter to be monitored. To conclude, the Workload corresponding to the AerT did not result statistically different between the two evaluation methods and showed in the ICC an excellent value of 0.95 [34–35]. However, this parameter showed a TE slightly higher than the acceptable limit (10.47%) [33].

These first results about the validity of the RQA method for the detection of the AerT could be summarized stating that the AerT_RQA_ is a valid method to use as alternative of gas analysis. In particular, the heart rate at AerT is considered the most valid parameter to monitor using the RQA method, and the determination of the AerT with this approach loses accuracy with VO_2_ and Workload variables. Between the different exercise variables, the HR seems to be not only the most accurate but also the most sustainable, indeed this parameter can be easily recorded during training using non-invasive, not expensive, time-efficient devices which can be applied routinely and simultaneously in a large number of athletes [13]. In the field of exercise assessment, the heart rate variability has been largely used to identify the AerT in several specific populations as diabetic and cardiac patients [37–38], healthy individuals [39–40], competitive cyclists and triathletes [41], and professional soccer players [11]. However, little is known about the accuracy of the alternative approach of the RQA of HRV time series. Indeed, this method so far has been used only for the identification of the AerT in obese subjects [2]. The present work enlarges the boundaries of this methodology confirming the efficacy of RQA analysis also in individuals with a higher physical fitness level. Therefore, the utilization of the AerT_RQA_ may provide an important training intensity guidance for the identification of the exercise zone which can be covered predominantly by the aerobic metabolism [4, 5, 17].

In terms of degree of agreement between AnT_GE_ and AnT_RQA_, all parameters (HR, VO_2_ and Workload) showed very strong correlations [36] and neither fixed or proportional biases were detected. In addition, mean percentage differences were less than 1% for HR and less than 2% for Workload and VO_2_ and no significant differences were found in all variables. Moreover, the acceptable TE values (HR 2.61%; VO_2_ 4.78%, Workload 6.64%) and the excellent ICC values (HR 0.89; VO_2_ 0.97; Workload 0.97) confirmed the validity of all parameters corresponding at the AnT_RQA_ [33–36]. As it was observed in the AerT, the lowest TE found in HR demonstrated that this parameter is the most accurate to use for the detection of the AnT with the RQA method.

Based on the aforementioned findings we believe that the RQA approach is a valid method as an alternative to gas analysis in the determination of the AnT. In addition, comparing the accuracy of the different variables, the heart rate at the AnT seems to be the most valid and sustainable parameter to use with the RQA method. The HRV has been utilized to estimate the AnT in different exercise evaluation fields [7–12, 38, 41–42]. However, to our knowledge, this is the first study which attempted to directly examine the non-linear approach of the RQA of HRV time series in the estimation of the AnT. Considering the high importance that the AnT detection has for sport performance [5, 7–8], these results may provide a useful contribution in training prescription and evaluation.

Even though we suggested the use of RQA method as a COVID-safe alternative to gas analysis for the detection of both metabolic thresholds, comparing the validity of the AerT_R_QA and AnT_RQA_ (with OLP regression, TE, ICC and the Bland Altman plots) it appeared that the AnT_RQA_ has a better accuracy than AerT_RQA_ and other studies showed similar results [9–10, 43]. This lower accuracy in the AerT_RQA_ may be explained by methodological problems in the determination of this threshold using gas analyzers. Indeed, it seems that the accuracy, the intra- and inter-observer reliability, the repeatability and the percentage of indeterminate thresholds of the AerT_GE_ had been questioned [5, 43]. Therefore, the lower agreement between AerT_RQA_ and AerT_GE_ may be caused by methodological problems on the GE method in some particular individuals. In view of all these aspects, the use of the RQA method could be used as a supportive approach to gas analysis for all the cases in which the detection of the AerT is ambiguous.

The following limitations are acknowledged. First, the number of participants in the study was relatively small. Second, the population was not heterogeneous (composed by 28 males and only 2 females). Thus, considering that metabolic thresholds are influenced by phenotypical sex differences [44], further research should investigate the RQA of HRV time series separately in males and females. In addition, future physiological exercise studies should assess if the findings of the present work could be adopted in particular field tests which do not necessarily require the use of a cycle-ergometer.

## Conclusion

The recurrence quantification analysis of heart rate variability time series is a COVID-safe approach for the determination of metabolic thresholds in individuals with different physical fitness levels, therefore, it can be used as a valid method for threshold detection alternative to gas analysis. Thus, in on-field evaluation, during an incremental exercise test it is possible to assess sport performance and delineate the intensity of training zones using non-invasive and low-cost devices. Based on the present results, coaches could obtain information about the training trend of their athletes avoiding classical respiratory measurement devices which require specific knowledge and that may be not allowed during this particular pandemic period.

## List of abbreviations

HR: Heart Rate
HRV: Heart Rate Variability
CPET: Cardiopulmonary exercise test
RPE: Rate of Perceived Exertion
RQA: Recurrence Quantification Analysis
GE: Gas Exchange
AerT_GE_: Aerobic Gas Exchange Threshold
AnT_GE_: Anaerobic Gas Exchange Threshold
AerT_RQA_: Aerobic Recurrence Quantification Analysis Threshold
AnT_RQA_: Anaerobic Recurrence Quantification Analysis Threshold
DET: percent of determinism, the percent of recurrence points which form diagonal lines
LAM: percent of laminarity, the percent of recurrence points which form vertical lines
BMI: Body Mass Index
BIA: Bioelectrical Impedance Analysis
FM%: Fat Mass percentage
VO_2_: Volume of Oxygen consumption
VCO_2_: Volume of Carbon dioxide production
VE: ventilation
OLP: Ordinary Least Product
TE: Typical percentage Error
ICC: Intraclass correlation coefficients
IC: confidence interval

